## Supplementary Figures for "A Model of Hippocampal Replay Driven by Experience and Environmental Structure Facilitates Spatial Learning"

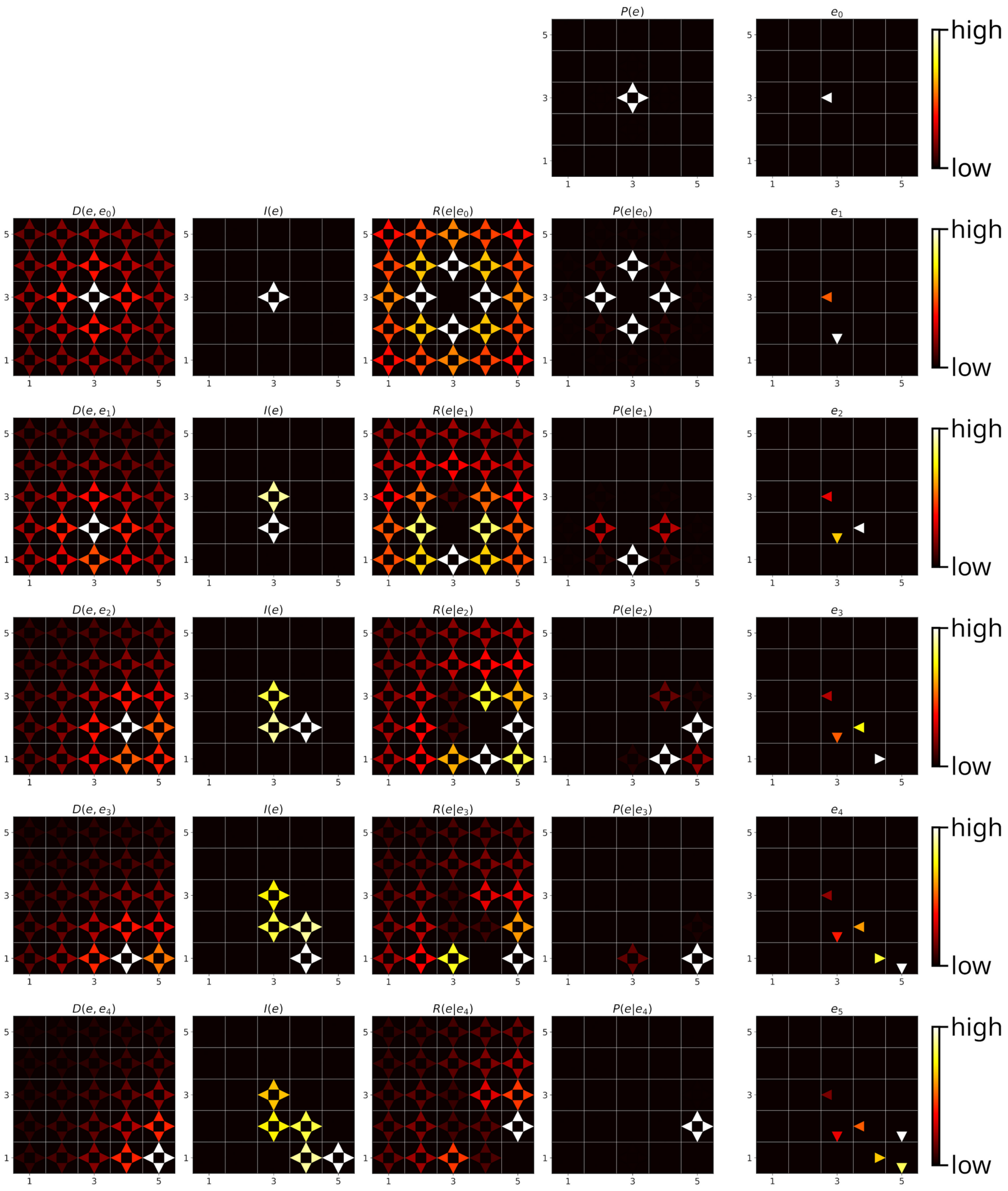


**Supplementary Figure 1: Step by step example of a replay sequence generated by SFMA.** From left to right: the different variables of SFMA as well as the reactivated experiences. Experience strength C(e) had the same value for all experiences and was therefore omitted. From top to bottom: the replay steps from start to finish. Replay was initiated at the center of the environment. Note that for replay initialization the prioritization variables do not convey information and were therefore omitted.

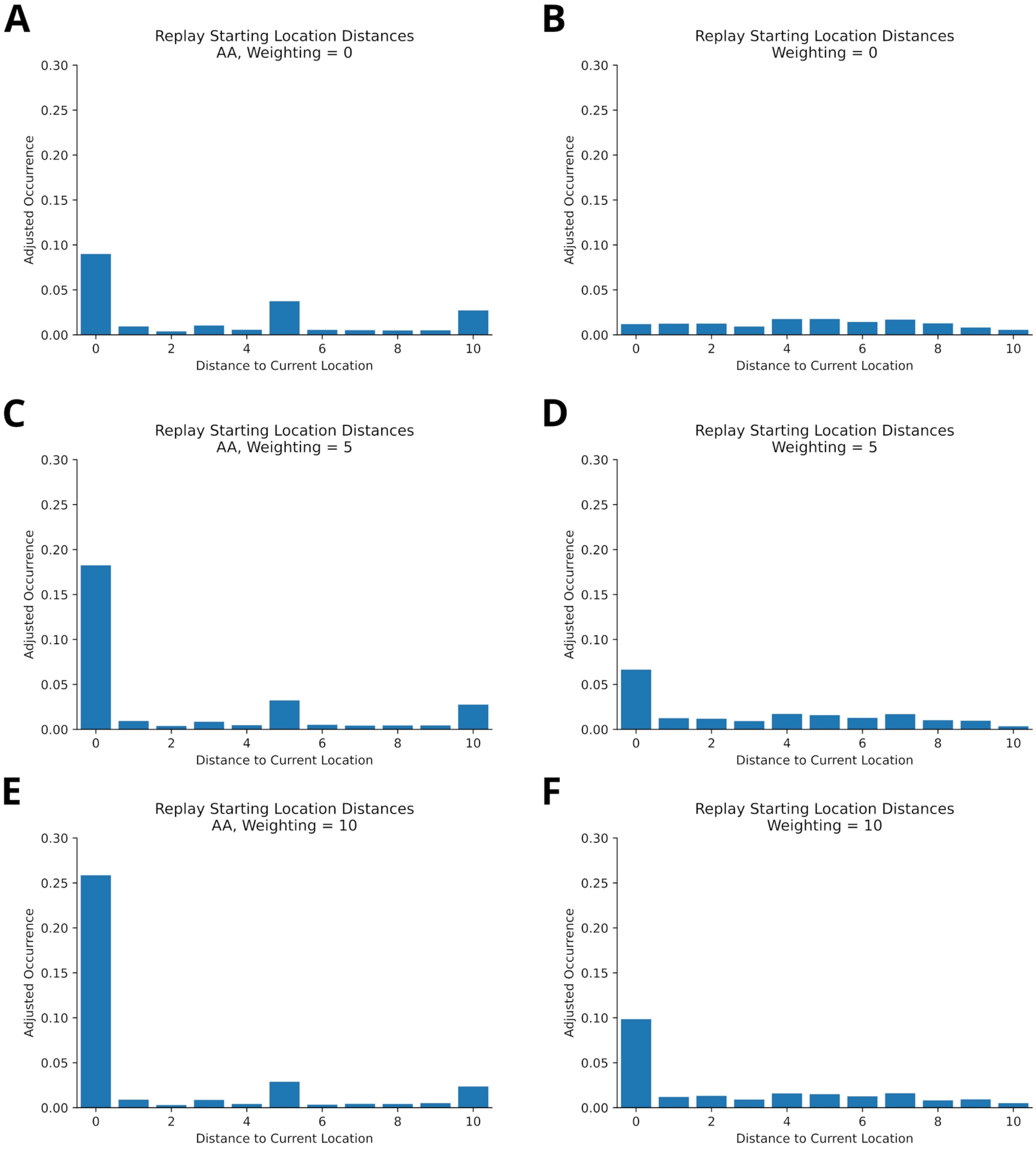


**Supplementary Figure 2: Reconciling non-local replays and preferential replay for current location in online replay. A**: Histogram of starting locations of *offline* replay recorded in the virtual version of Gupta et al.'s experiment. Raw occurrences for each bin were divided by the maximum number of bin occurrences to account for uneven distribution of distance bins. Replays were recorded at either of the reward locations. Current locations are over-represented, but replay is also initiated at other locations (non-local replay). **B**: Same as in A, but in an open field environment with no reward and homogeneous experience strengths. Replays were recorded at different locations in the environment. Non-local replays are prevalent, but there is no preference for the current location – contrary to observations of *online* replay. **C/D**: Same as A/B, but the experience strengths close to current location were increased to model an initialization bias. In both environments, the current location is more strongly represented by replays and non-local replays occur. **E/F**: Same as C/D, but with a stronger increase of experience strengths. The current location is more even more strongly represented by replays.

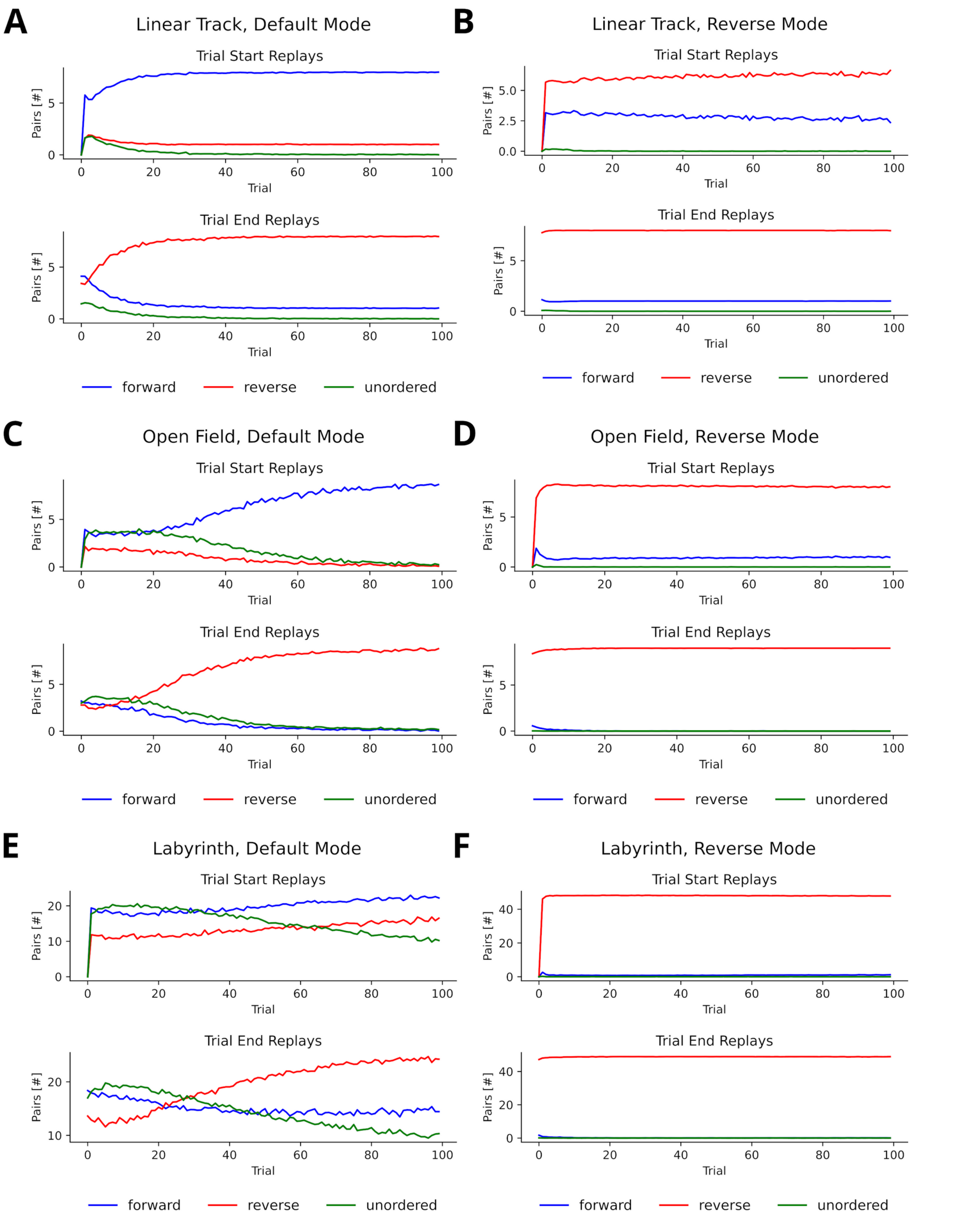


**Supplementary Figure 3: Directionality of consecutive replay pairs. A**: Pairs produced on a linear track when using the default mode for replays generated at the start (top) and end (bottom) of a trial. A pair of consecutively reactivated experiences $e_{t-1}$ and $e_t$ was considered forward when the next state of $e_{t-1}$ was the current state of $e_t$, reverse when the current state of $e_{t-1}$ was the next state of $e_t$ and unordered otherwise. **B**: Pairs produced on a linear track when using the reverse mode. **C/D**: Same as A/B, but in an open field environment. **E/F**: Same as A/B, but in a labyrinth environment.

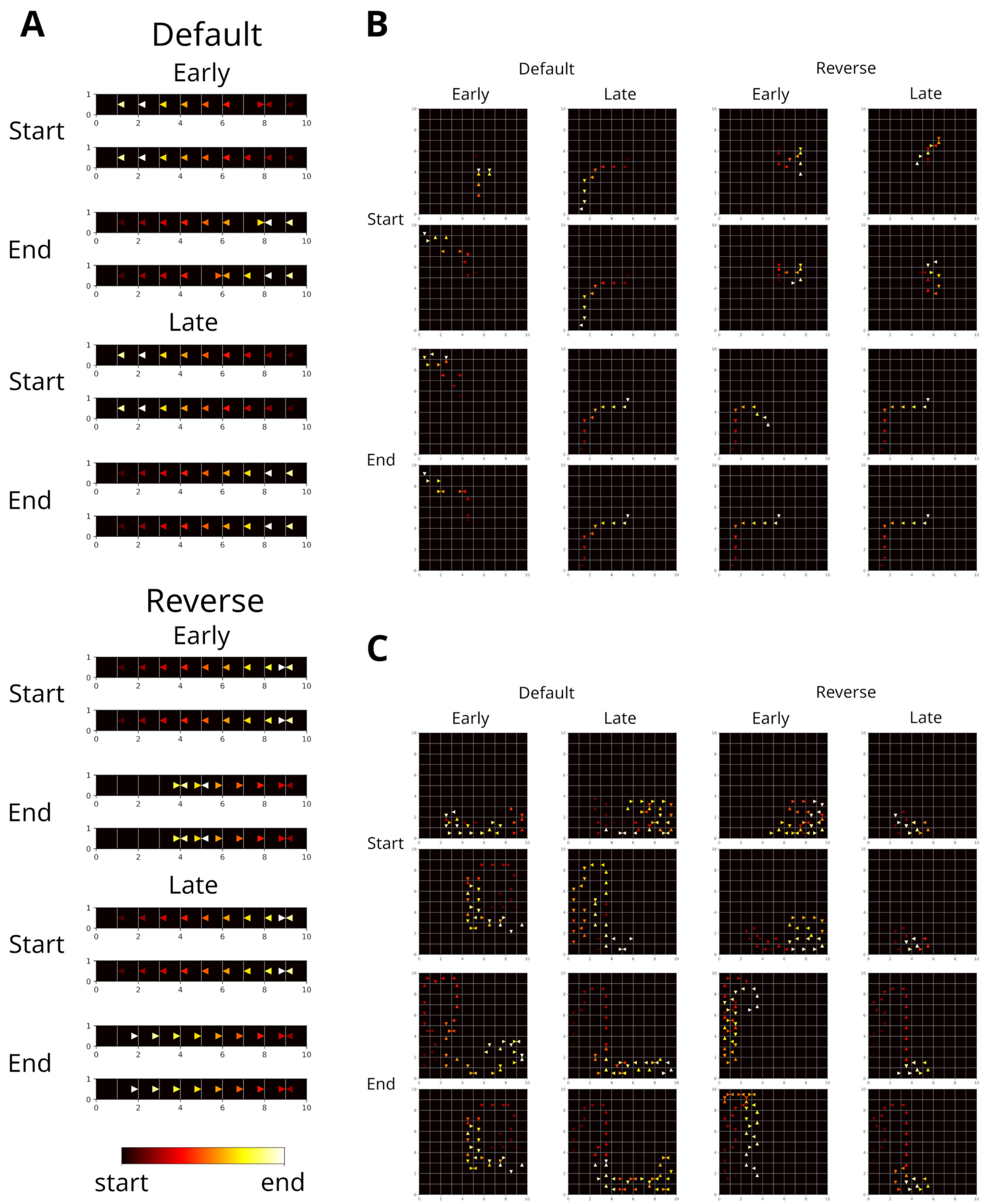


**Supplementary Figure 4: Example replay trajectories in different spatial navigation tasks.** **A**: Replay trajectories generated at the start and end of trials in the linear track for different replay modes (default and reverse) and different stages of learning (early and late). Color indicates when in a replay event a transition was replayed. **B**: Same as A, but in an open field. **C**: Same as A, but in a labyrinth like in Widloski and Foster (2022).

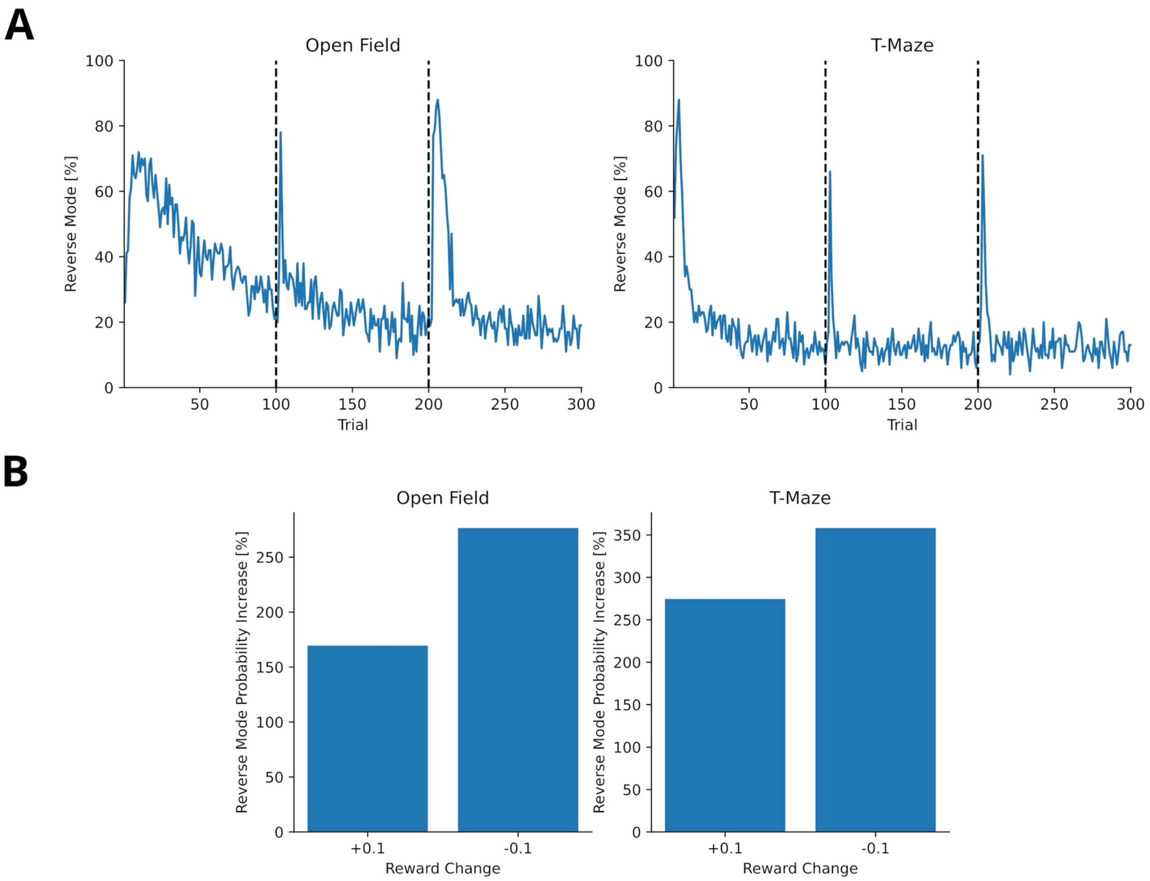


**Supplementary Figure 5: Reward changes trigger higher probability of activating the reverse mode when using the dynamic mode. A**: The probability of replay being generated in the reverse mode for agents trained in open field (left) and T-maze (right) for 300 trials. Reward was changed once after 100 trials (+0.1) and again after another 100 trials (-0.1). The probability for reverse mode spikes once in the beginning when the reward was first encountered and when the reward changes. **B**: The percentage change of reverse mode activation after reward change. Percentage change was computed from the five trial average before and after the reward change.

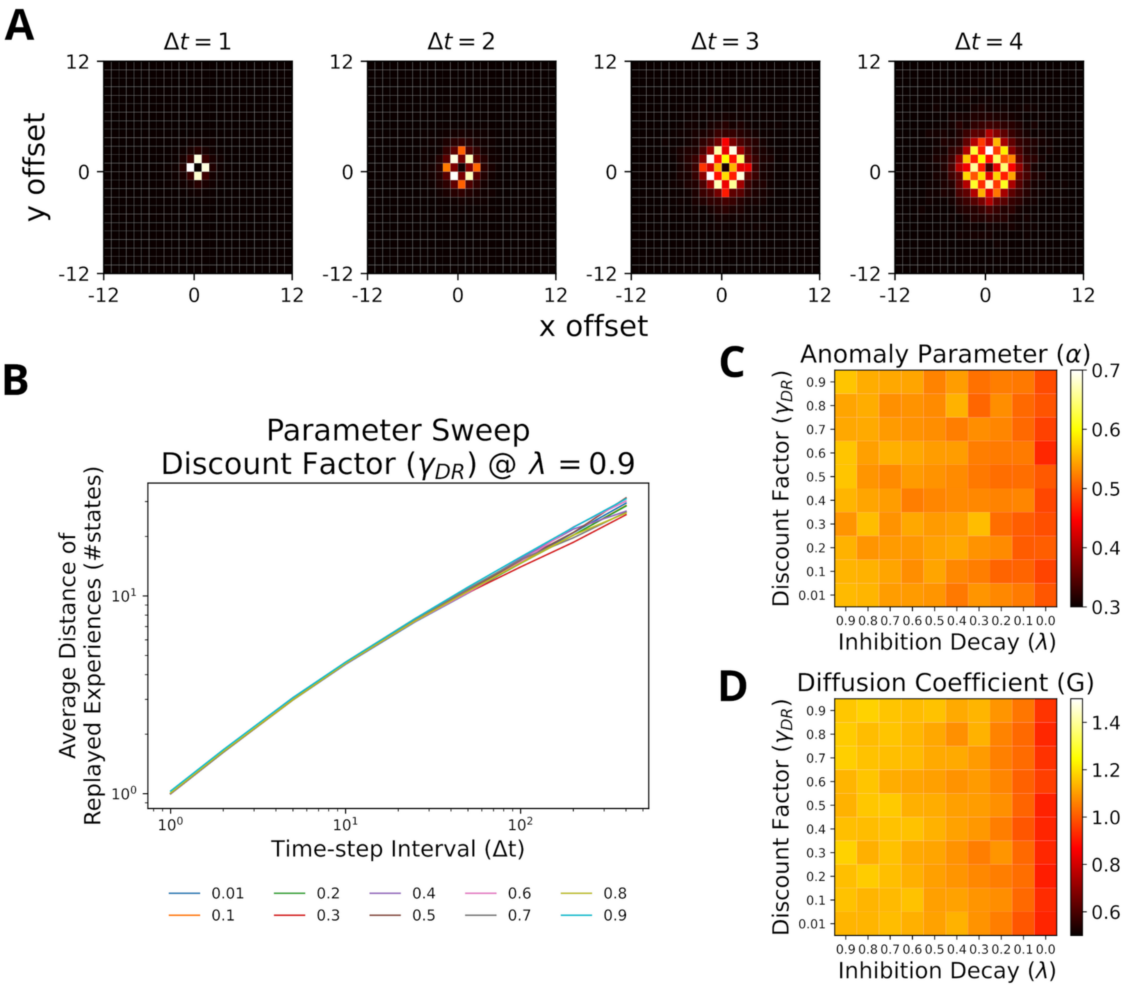


**Supplementary Figure 6: Replays generated using the reverse mode also resemble random walks across different parameter values for Default Representation (DR) discount factor and inhibition decay. A**: Displacement distribution for the first four time steps. **B**: Our model can reproduce the linear log-log relationship between time-step interval and average distance of replayed experiences. **C**: The anomaly parameters for different parameter values of DR and inhibition decay. Faster decay of inhibition yields anomaly parameters which closer resemble a Brownian diffusion process (i.e., closer to 0.5). **D**: The diffusion coefficients for different parameter values of DR and inhibition decay. Slower decay of inhibition yields higher diffusion coefficients.

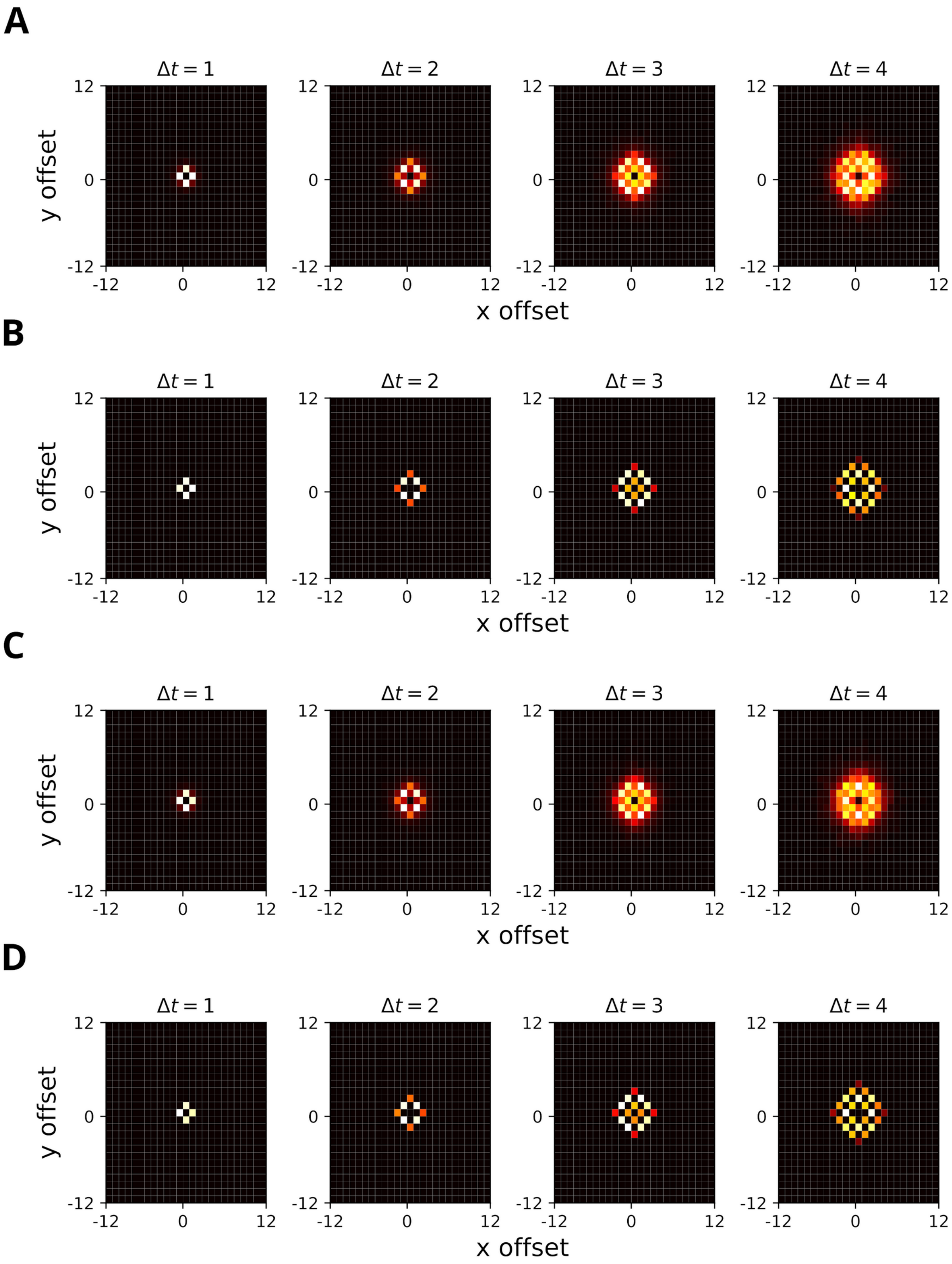


**Supplementary Figure 7: Displacement distributions for different inverse temperature values ($\gamma_{DR}$=0.9 and λ=0.9). A**: Displacement distribution for the first four time steps with $\beta_M$=5 given homogeneous experience strength. **B**: The same as A, but with a higher inverse temperature $\beta_M$=15. Note the lower local variance of the distribution compared to A. **C**: The same as A, but given heterogeneous experience strengths. **D**: The same as C, but with $\beta_M$=15. Note the negligible difference between homogeneous and heterogeneous experience strengths.

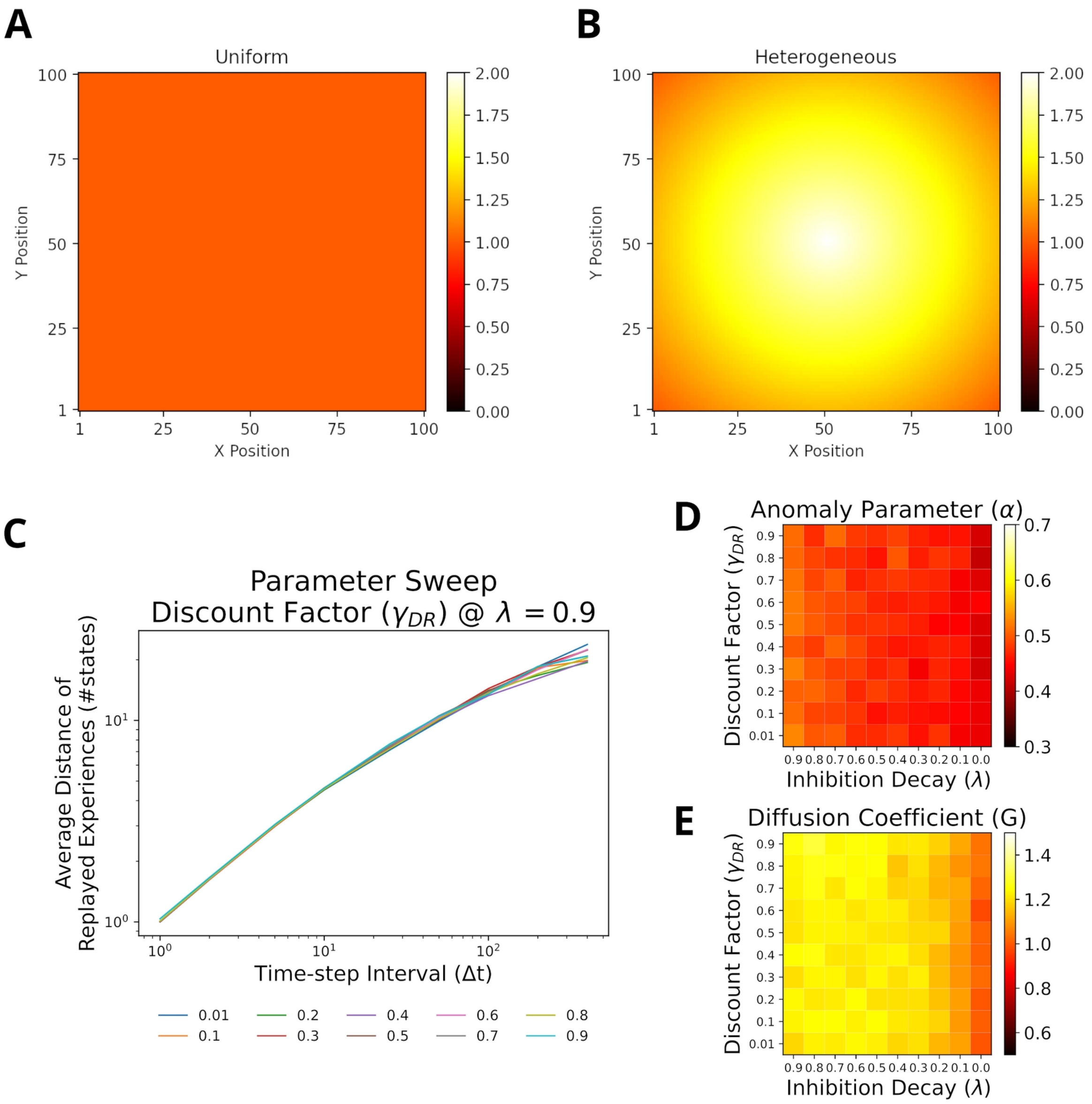


**Supplementary Figure 8: For heterogeneous experience strengths replays also resemble random walks across different parameter values for Default Representation (DR) discount factor and inhibition decay. A**: Homogeneous experience strength condition. **B**: Heterogeneous experience strength condition. **C**: Our model can reproduce the linear log-log relationship between time-step interval and average distance of replayed experiences. **D**: The anomaly parameters for different parameter values of DR and inhibition decay. Faster decay of inhibition yields anomaly parameters which closer resemble a Brownian diffusion process (i.e., closer to 0.5). **E**: The diffusion coefficients for different parameter values of DR and inhibition decay. Slower decay of inhibition yields higher diffusion coefficients.

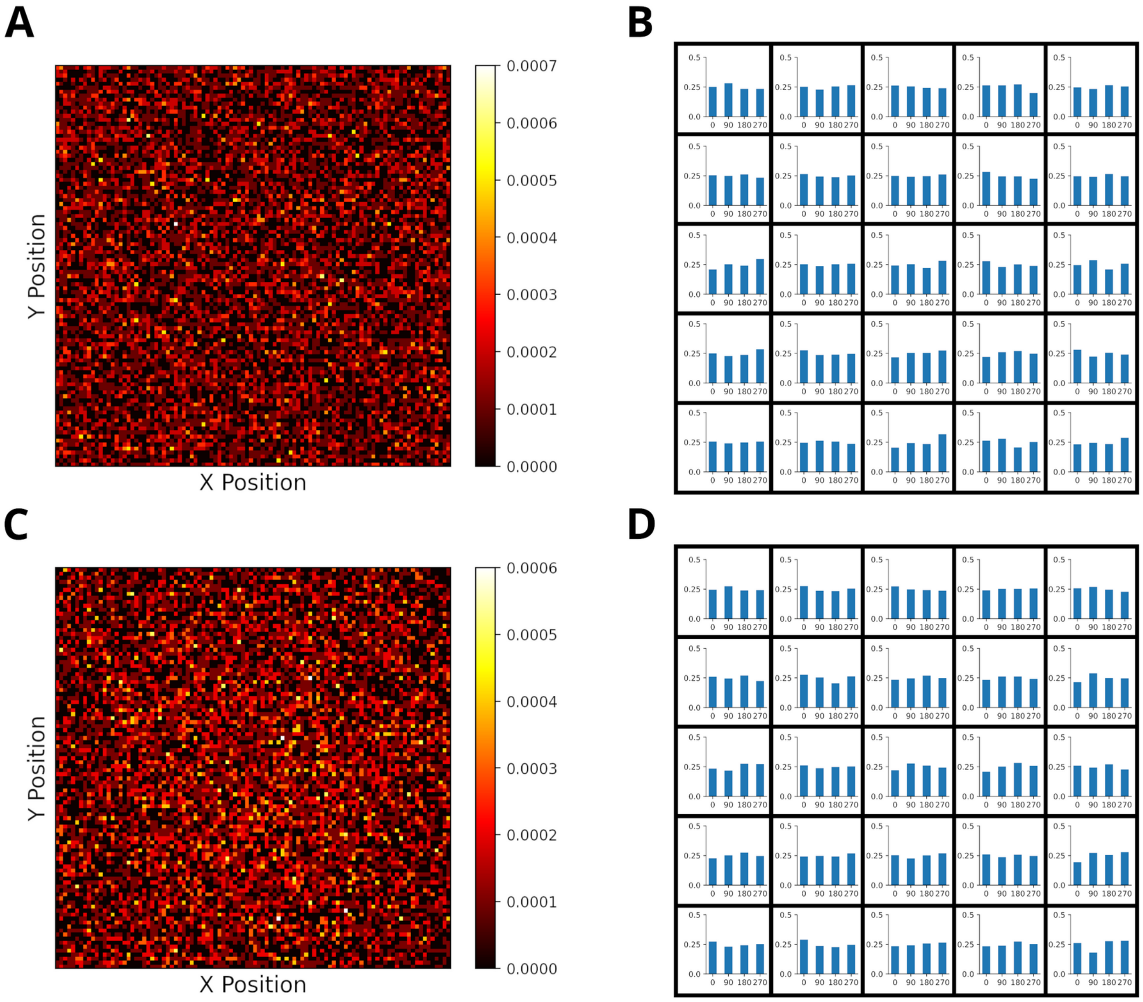


**Supplementary Figure 9: The starting positions of replay are randomly distributed across the environment in a 2-d open field. A**: Distribution of replay starting positions given homogeneous experience strengths. Starting positions are distributed (>90% randomness) evenly across the environment. Randomness of starting locations was measured like in Stella et al. (2019). **B**: Initial direction of replays are evenly distributed given homogeneous experience strengths. The environment was divided into bins of size 20x20. **C/D**: Same as A/B, but with heterogeneous experience strengths.

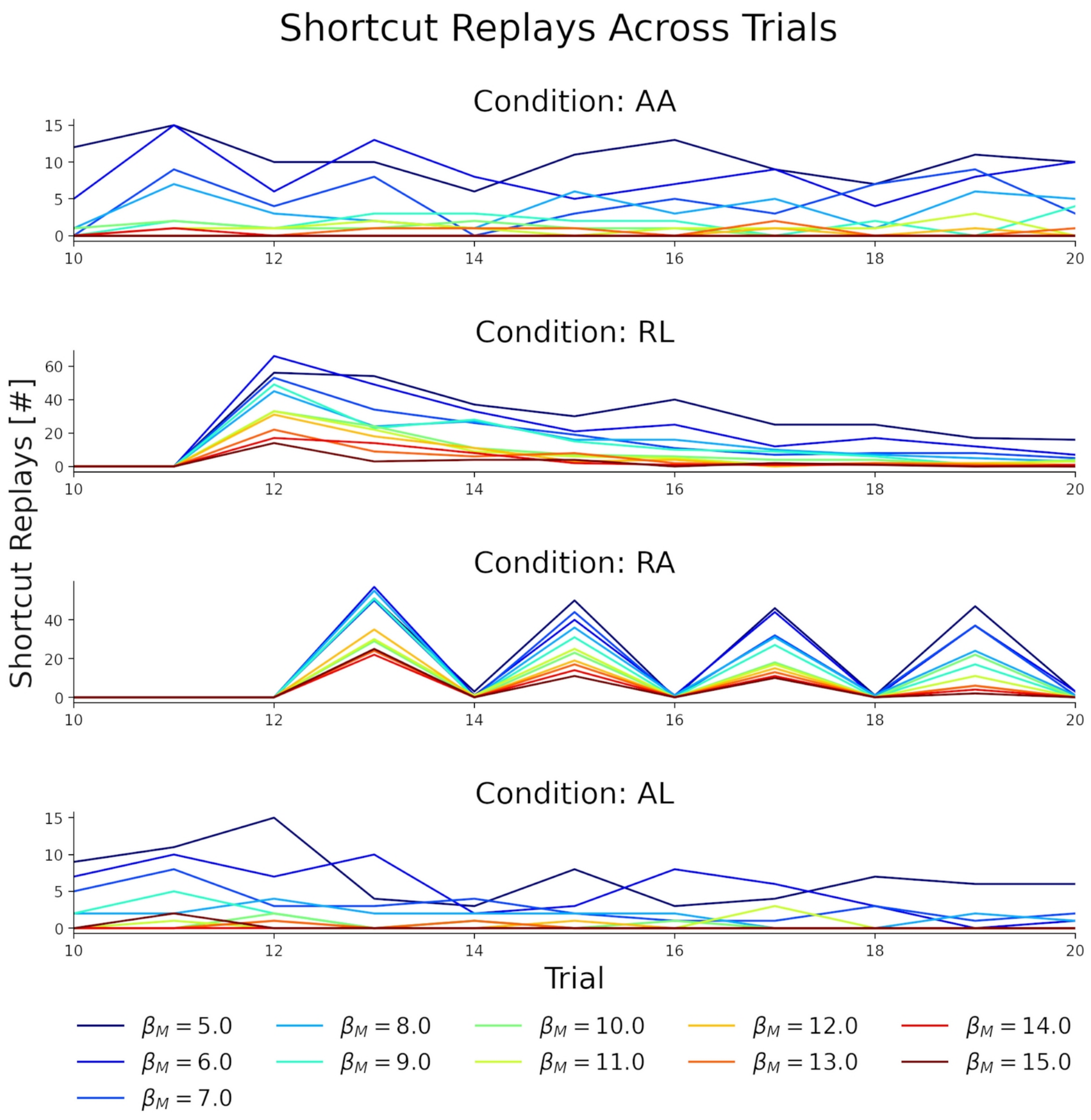


**Supplementary Figure 10: Number of shortcut replays on a trial by trial basis differs depending on behavioral statistics.** Shortcut replays in each trial for different experimental conditions: alternating-alternating (AA), right-left (RL), right-alternating (RA) and alternating-left (AL). For all conditions the number of shortcut replays is affected by the choice of the inverse temperature parameter.

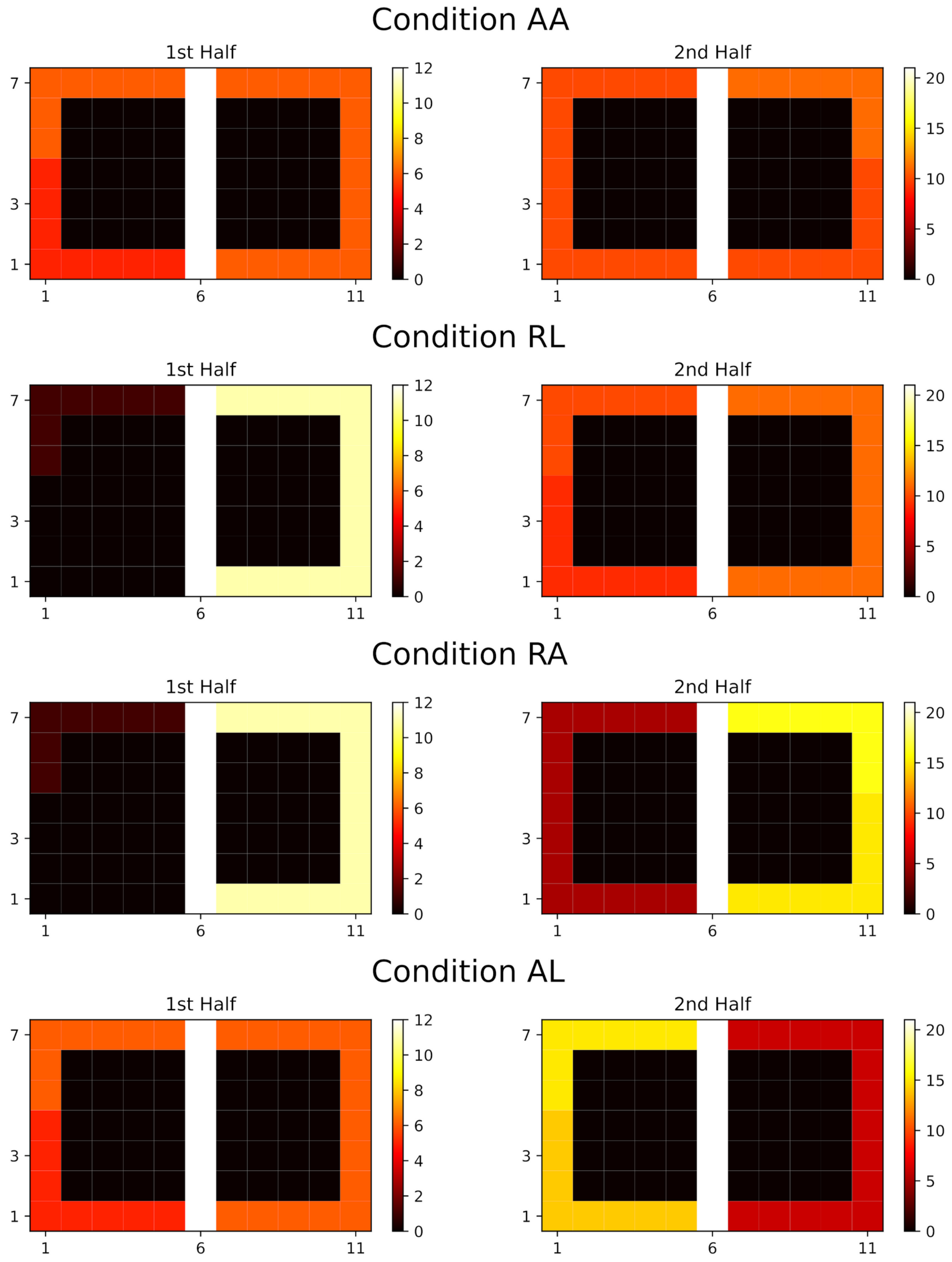


**Supplementary Figure 11: Experience strengths resulting from different behavioral statistics in Gupta et al.'s (2010) experiment.** Experience strengths after the first (left panels) and second half (right panels) given the different behavioral statistics (rows). For simplicity, the experience strength shown in each state is the sum over the four potential actions. Conditions: alternating-alternating (AA), right-left (RL), right-alternating (RA), and alternating-left (AL).

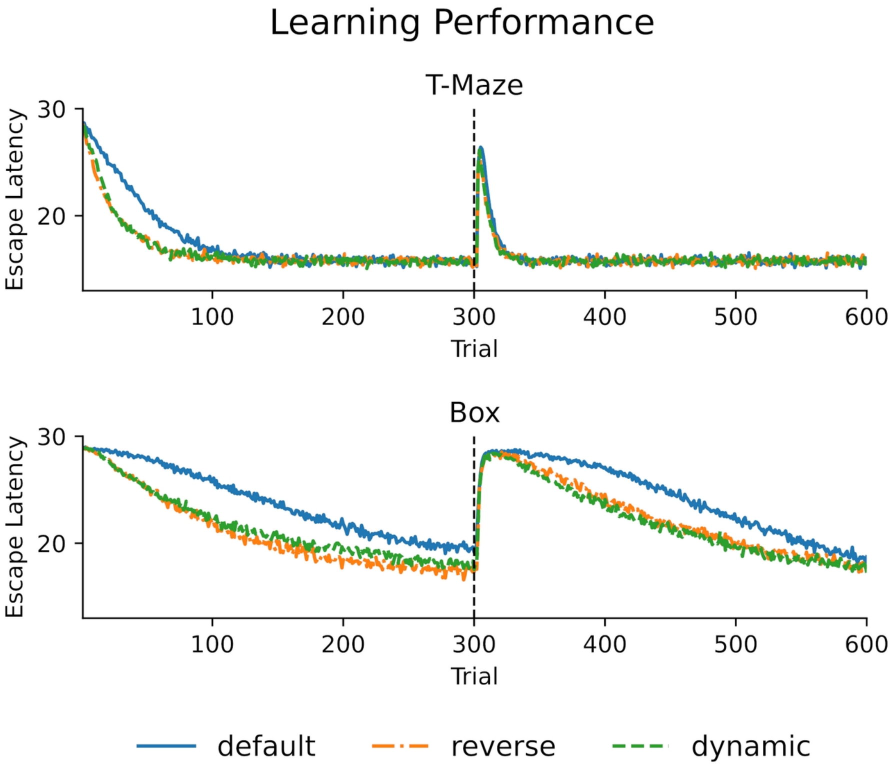


**Supplementary Figure 12: Following strongly stereotypical behavior efficient learning occurs for the default mode after a change in goal location. Top**: The learning performance for different replay modes in a T-maze environment. The goal arm changes after 300 trials. For learning of the initial goal the default mode is clearly outperformed by reverse and dynamic modes. However, there is no difference in learning performance between the different modes for learning the new goal location. **Bottom**: Same as for the top panel, but for an open field (Box), which allows for more variability in behavior (i.e., less stereotypical behavior). Learning performance is overall worse for all modes. The default mode is outperformed by the other two modes for both goal locations.

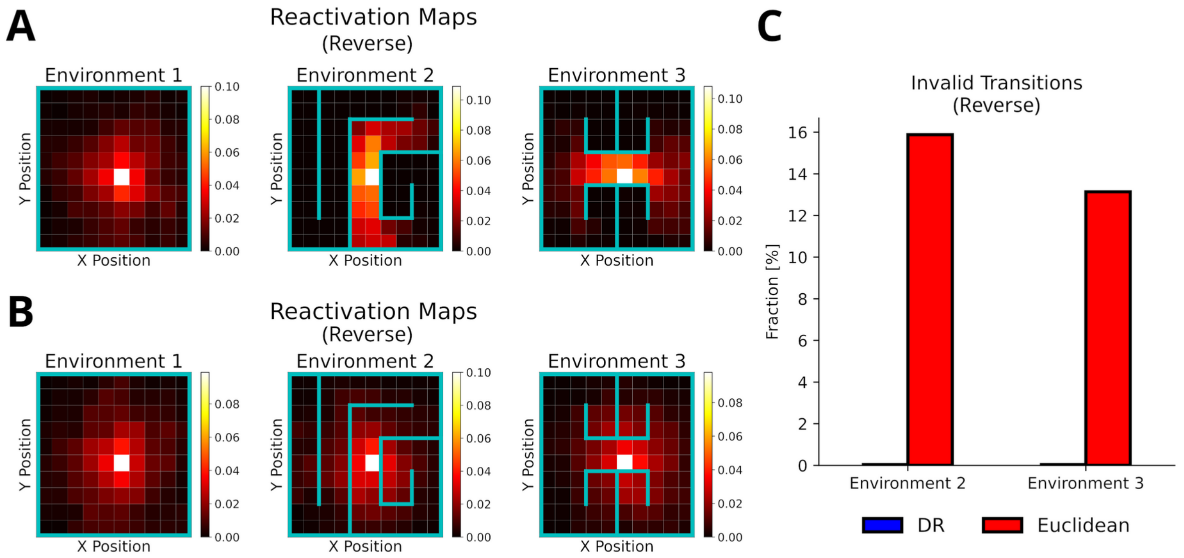


**Supplementary Figure 13: Reverse mode replays adapt to environmental changes. A**: Reactivation maps, i.e., the fraction of state reactivations, for reverse mode replay recorded in three environments ($\gamma_{DR}$=0.1). The color bar indicates the fraction of reactivations across all replays. Note that replay obeys the current environmental boundaries. **B**: Same as in A, but experience similarity is based on the Euclidean distance between states. Note that replay ignores the boundaries. **C**: The fraction of invalid transitions during reverse model replay in different environments for the DR (blue) and Euclidean distance (red). While replay can adapt to environmental changes when using the DR, it does not when experience similarity is based on the Euclidean distance.

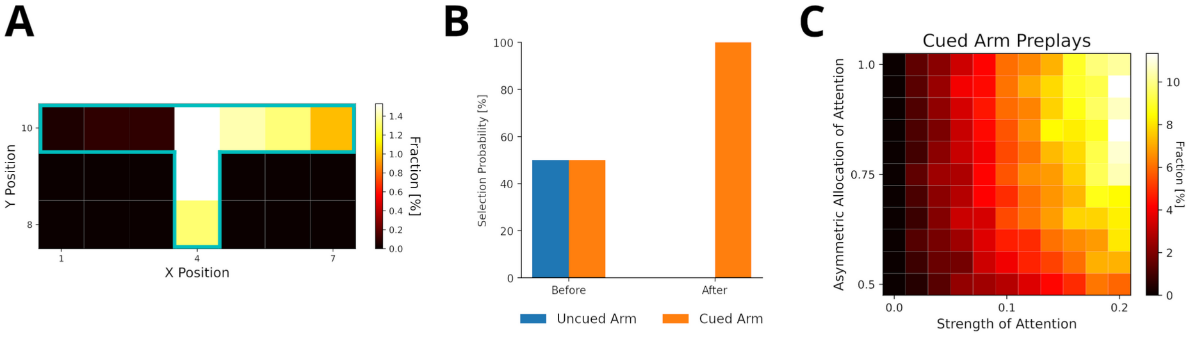


**Supplementary Figure 14: Preplay of yet unvisited cued locations can be explained by visual exploration (reverse mode). A**: Fractions of reactivated arm locations. Experiences associated with the cued arm (right) are preferentially reactivated. **B**: The fractions of choosing the cued arm and uncued arm before and after training the agent with preplay. The agent preferentially selects the cued arm after preplay. **C**: The percentage of cued-arm preplay for different amounts of attention paid to the cued arm vs. the uncued arm.

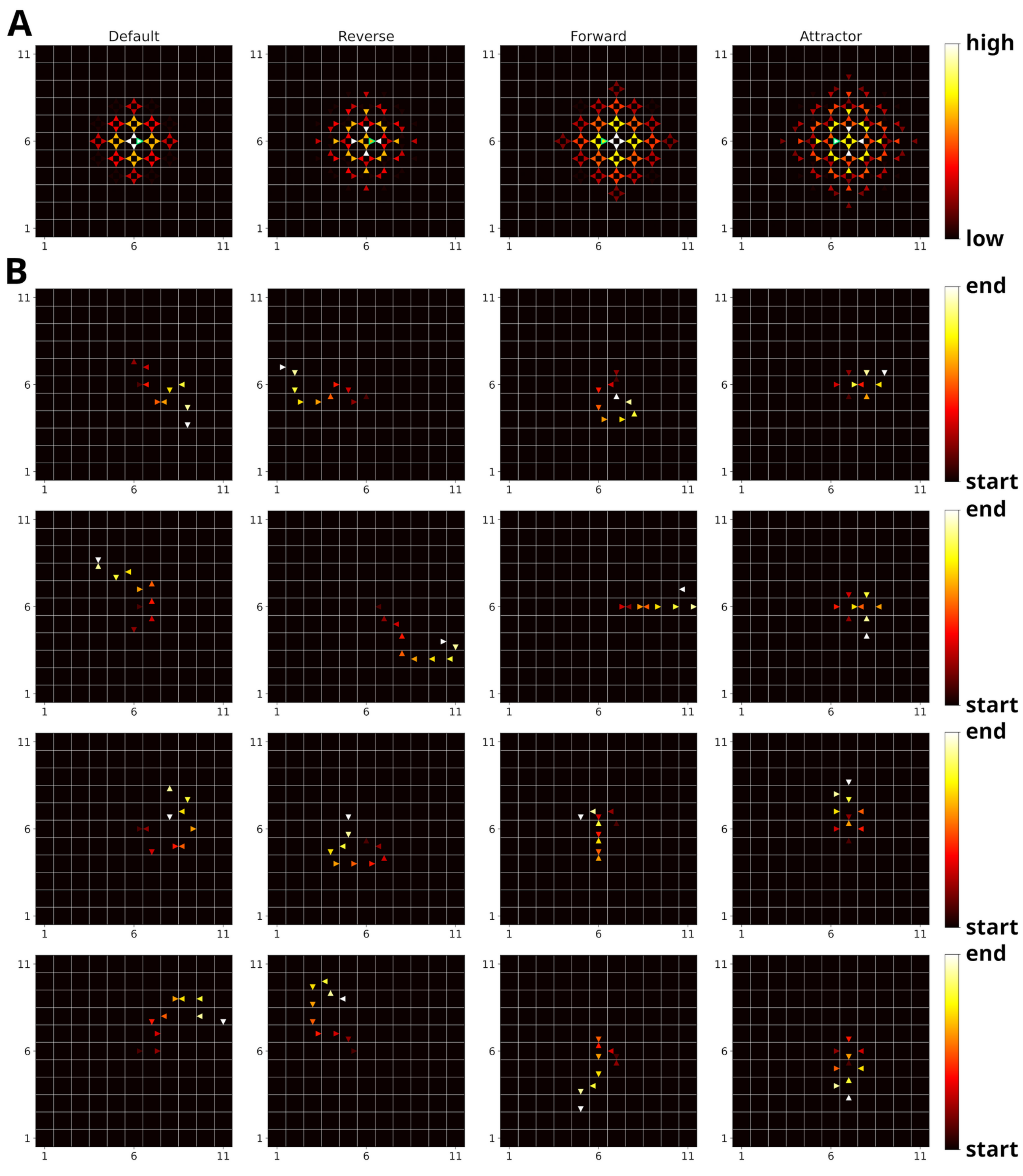


**Supplementary Figure 15: Possible replay modes and example replay trajectories. A**: Experience similarities of the different replay modes, i.e., default, reverse, forward and attractor mode, for a given experience (marked in green). **B**: Example replay trajectories generated in an open field environment with the different replay modes (from left to right: default, reverse, forward and attractor mode). Experience strengths were homogeneous and replay was initiated in the center of the environment.

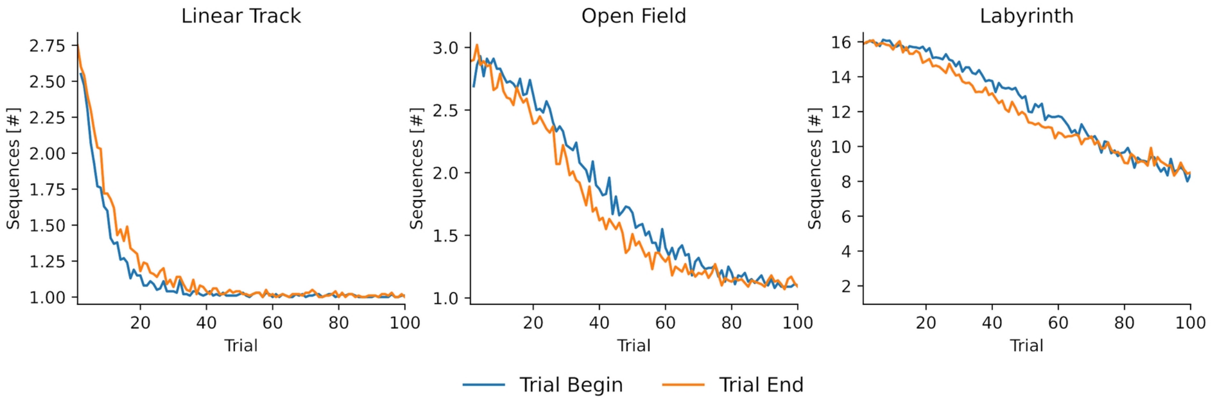


**Supplementary Figure 16: The number of sequences decreases with experience.** The number of sequences per replay as a function of experienced trials for three different environments, i.e., linear track, open field and labyrinth. The replay lengths were 10, 10 and 50, respectively. Early during learning replay contains multiple short sequences. By the end of learning replay tends to represent one long sequence in linear track and open field environments. For the labyrinth environment the number of sequences is still decreasing due to the complexity of the environment and the much higher replay length.

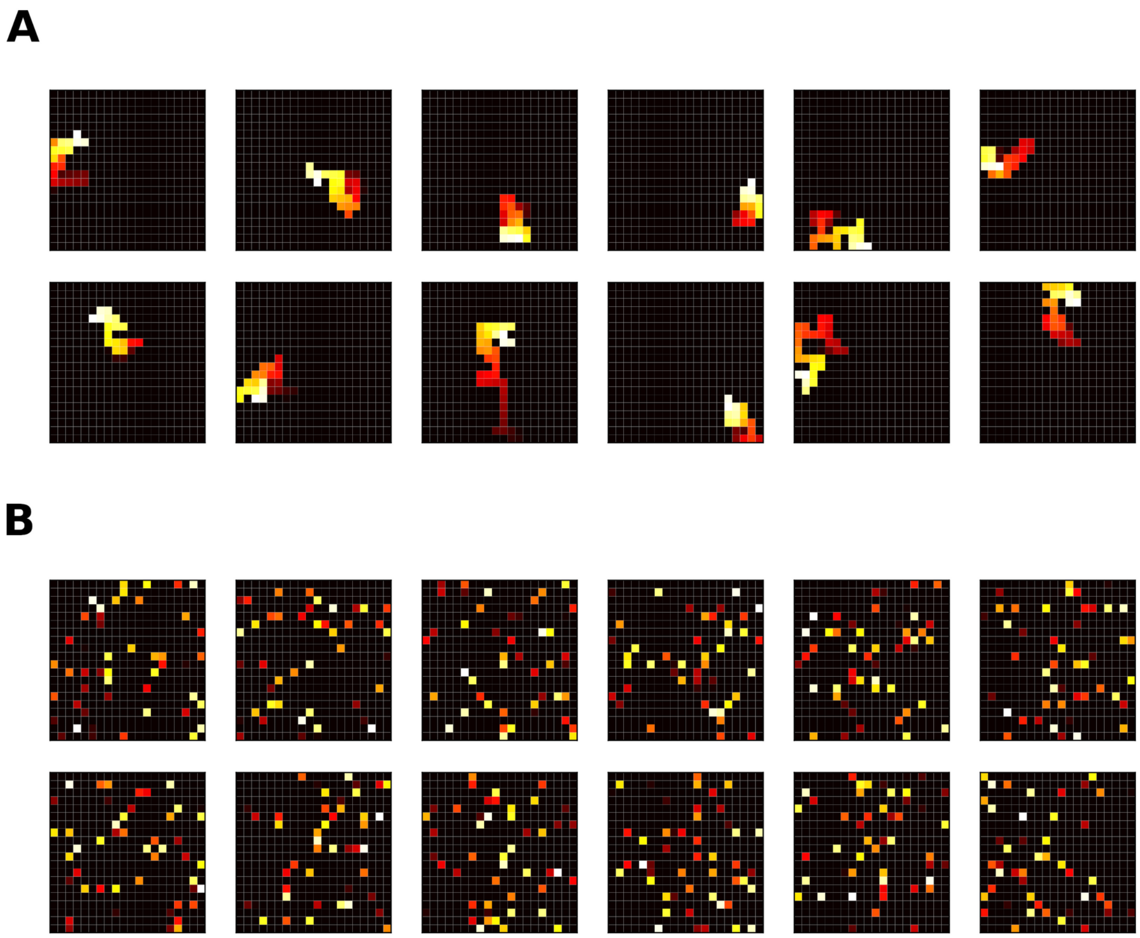


**Supplementary Figure 17: Prioritized Memory Access' (PMA) ability to produce sequences is severely disrupted when the gain calculation for n-step updates is adjusted. A**: Sequences generated by PMA (Mattar and Daw, 2018) given uniform need and all-zero gain. Because utility is defined as a product of gain and need, the gain must have a nonzero value to prevent all-zero utility values. This is achieved by applying a (small) minimum gain value. For n-step updates, which update the Q-function for all n steps along the trajectory and are the main driver for forward replay sequences in PMA, the gains are summed across all n steps. Mattar and Daw (2018) apply the minimum gain value before summation, which artificially increases the gain of n-step updates. **B**: Same as A, but the minimum gain value is enforced after summing gain along the steps of the n-step update. We argue that the minimum value should be applied after the summation of gain values, since otherwise the gain artificially increases with sequence length. If done so, PMA loses the ability to produce forward sequences and starts to randomly reactivate experiences.
